# Object-finding skill created by repeated reward experience

**DOI:** 10.1101/043786

**Authors:** Ali Ghazizadeh, Whitney Griggs, Okihide Hikosaka

## Abstract

For most animals, survival depends on rapid detection of rewarding objects, but search for an object surrounded by many others is known to be difficult and time consuming. However, there is neuronal evidence for robust and rapid differentiation of objects based on their reward history in primates (Hikosaka et al., 2014). We hypothesized that such robust coding should support efficient search for high-value objects, similar to a pop-out mechanism. To test this hypothesis, we let subjects (n=4, macaque monkeys) view a large number of complex objects with consistently biased rewards with variable training durations (1, 5 or >30days). Following training, subjects searched for a high-value object (Good) among a variable number of low-value objects (Bad). Consistent with our hypothesis, we found that Good objects were accurately and quickly targeted, often by a single and direct saccade with a very short latency (<200ms). The dependence of search times on display size reduced significantly with longer reward training, giving rise to a more efficient search (40ms/item to 16ms/item). This object-finding skill showed a large capacity for value-biased objects and was maintained in the long-term memory with no interference from reward learning with other objects. Such object-finding skill, particularly its large capacity and its long term retention, would be crucial for maximizing rewards and biological fitness throughout life where many objects are experienced continuously and/or intermittently.

**Significance Statement:** Visual objects that have become associated with reward in the past, can grab our attention even when we are not looking for them. Here, we show that this powerful attentional mechanism serves an important biological purpose: it allows one to quickly find valuable objects regardless of the number of other visual distractors present. Efficient search has long been thought to be primarily limited to objects with certain visually conspicuities (Wolfe and Horowitz, 2004). Our result shows that long-term and consistently biased reward can achieve search efficiencies that are independent of object visual features. This search efficiency is highly scalable as it develops for a large number of objects with no apparent interference between objects and is maintained in long-term memory.

## Introduction

Surrounded by many objects, animals often need to quickly find valuable objects, such as food. In humans and monkeys, object identification is best done at the fovea and degrades in the periphery (Low, 1951; Rentschler and Treutwein, 1985; Strasburger et al., 2011). Due to these perceptual/attentional limitations (Xu and Chun, 2009), search for a target object can be difficult (Wolfe and Bennett, 1997) and may require multiple shifts of gaze (saccades) (Zelinsky and Sheinberg, 1997; Motter and Belky, 1998). Yet, based on some recent findings, we speculated that such difficulty may be overcome by certain ecological experiences. Studies from our laboratory suggest that the caudal part of the basal ganglia which is involved in oculomotor and attentional control, differentially responds to high-and low-valued objects (Yasuda et al., 2012; Kim and Hikosaka, 2013; Yamamoto et al., 2013). In particular, substantia nigra reticulata (SNr) neurons projecting to the superior colliculus automatically and rapidly (<200ms) discriminate stably high-and low-valued objects (Yasuda et al., 2012). They have long-term memories of object values (> 100 days) with a high capacity (> 300 experienced objects). Behaviorally, monkeys were found to exhibit strong gaze bias toward objects with memory of high-reward when freely viewing multiple objects (Yasuda et al., 2012; Kim and Hikosaka, 2013; Yamamoto et al., 2013). These findings are consistent with studies on human subjects showing that previously reward-associated objects can automatically distract attention (Anderson et al., 2011; Chelazzi et al., 2012; Theeuwes and Belopolsky, 2012).

These results suggest that repeated reward experience should enable primates to efficiently locate Good objects when searching for reward. However, this hypothesis has not been directly tested, because in the previous studies, the subjects did not look for objects based on reward experience. Therefore, the following critical questions remain: How quickly and accurately does attention/gaze reach a Good object during search? How does reward history affect search efficiency when one is confronted with increasing number of distractors? To answer these questions, we used a visual search task. Instead of focusing on the effects of visual features, we examined how reward history of objects controls object search.

## Results

To study the effect of reward experience on visual search, we let monkeys (n=4) view many fractal objects (Figure 1A), only one in each trial (Figure 1B), and did so across multiple sessions (Figure 1D). Each object was consistently associated with a large or small reward (Good and Bad fractals) (Figure 1A). Following a given number of object-value learning sessions, we tested the monkey’s ability to find Good objects (target) using a search task (Figure 1C-D). A Good object was present in half of the trials (see Methods). Before the final choice, the monkey was allowed to make multiple saccades (Figure 1C, 2A).

**Figure 1.**
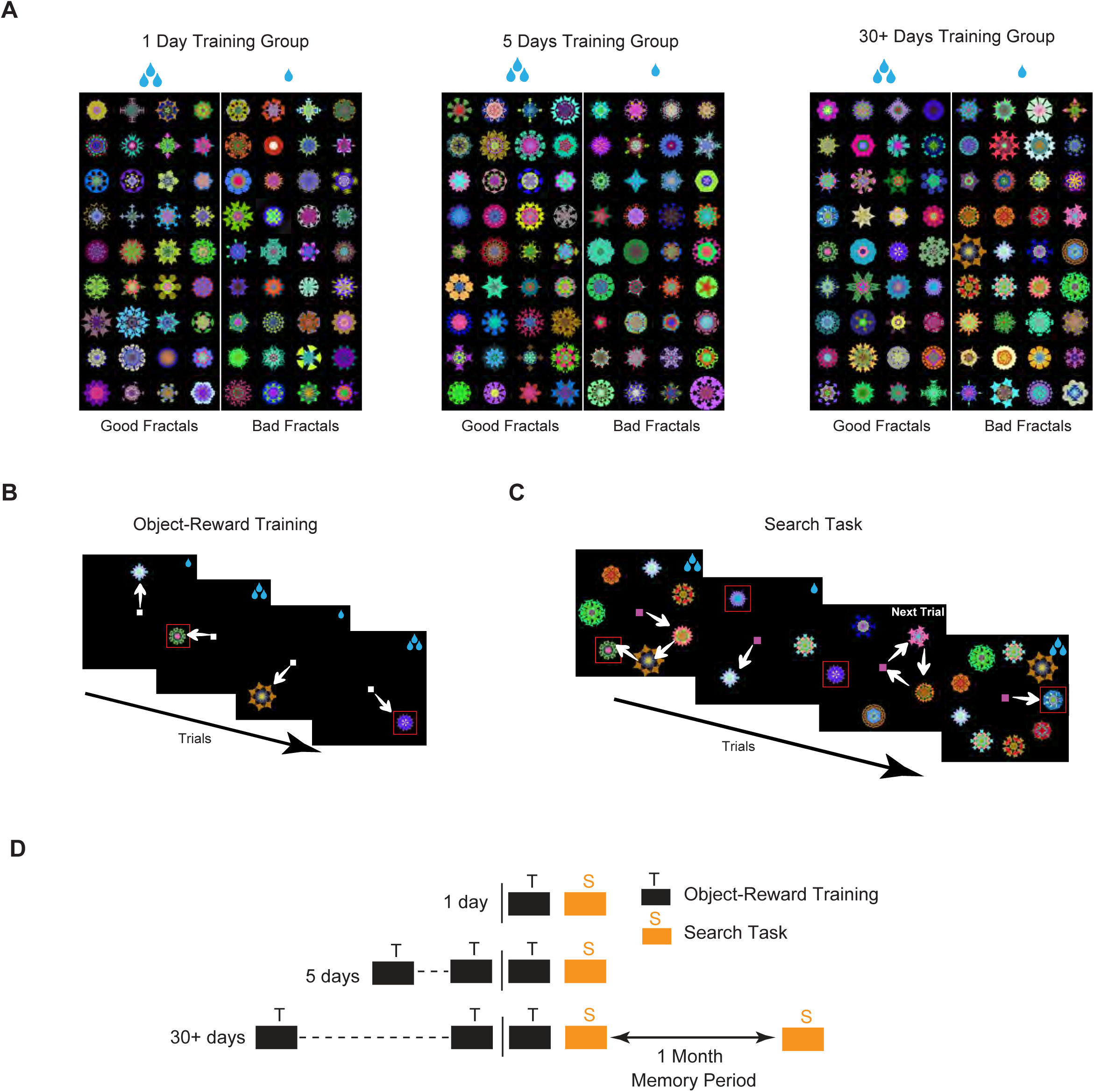
Training and testing procedures. **A,** Fractal objects for monkey R (n=212), each of which was consistently associated with a large reward (Good objects) or a small reward (Bad objects). **B,** Object-reward training: A large or small reward was delivered after the monkey made a saccade to a fractal object. The fractal was presented in one of 8 directions peripherally (15°). **C,** Search task (test): A large or small reward was delivered after the monkey chose one among multiple fractals. The choice was determined by gaze longer than 500ms so that the monkey was allowed to make multiple saccades with shorter gaze durations. Monkey could return to center to go to the next trial if no Good fractal was found. The number of simultaneously presented fractals varied across trials (15° eccentricity, display size = 3, 5, 7, 9). Among them, only one fractal was Good object in half of the trials. In the other half, no Good object was present (not shown). In **B** and **C,** red square indicates Good object (not shown to the monkey). **D,** Each monkey viewed three separate groups of reward-biased fractals (Figure 1A) with different training amounts (1, 5, or 30+ days, 72 fractals/group) followed on the last day (vertical line) by the search task (Figure 1C). Fractals in the ‘30+ days’ group were tested again after 1 month (Mem, in subsequent figures).

**Figure 2.**
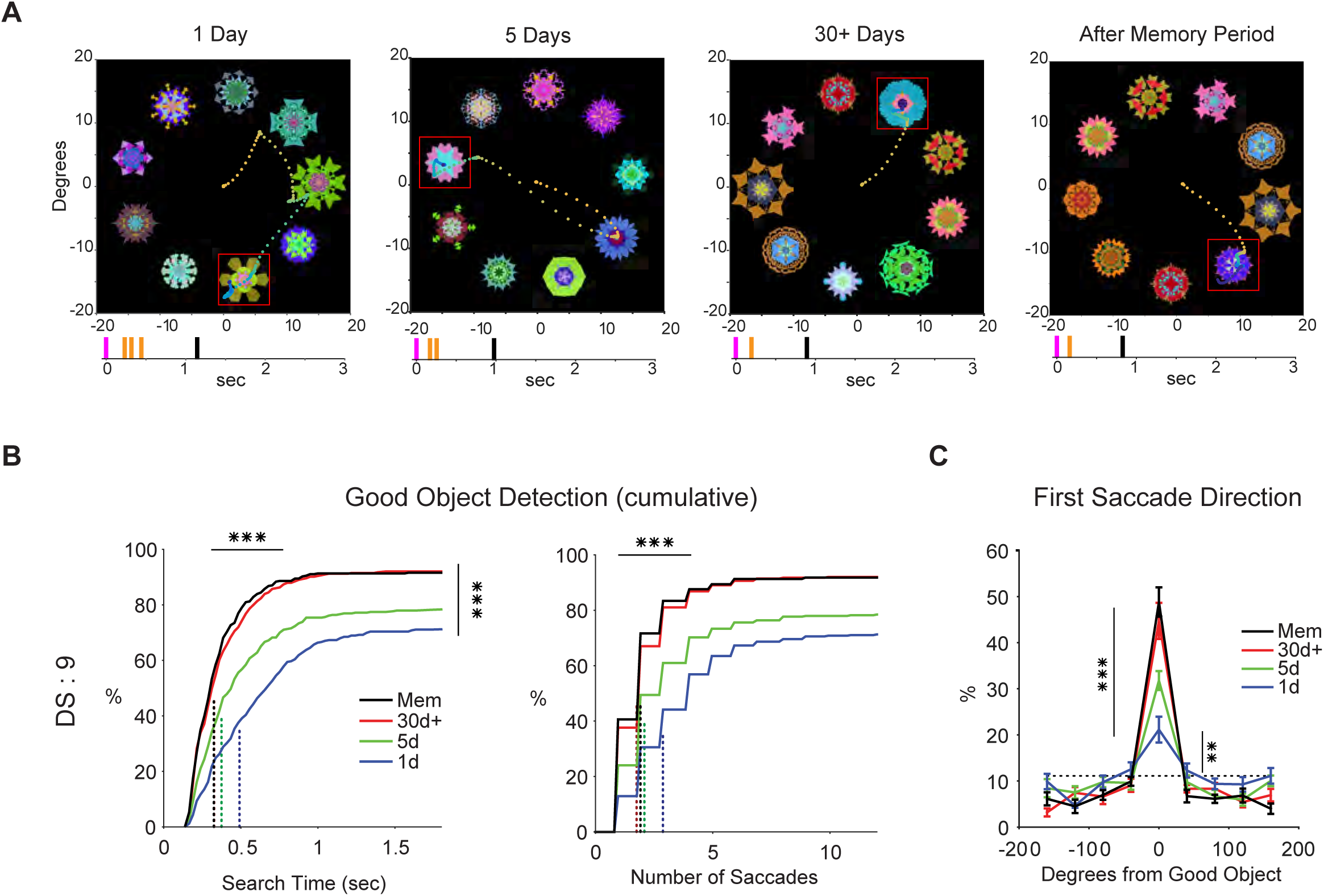
Effects of the repeated object-reward association on the detection of Good objects (DS: 9). **A,** Example search performance of monkey R after different training amounts and memory period. Eye position is shown by time-dependent color-coded dots (2ms/sample, from orange to blue). Red square indicates Good object (not shown to the monkey). Tick marks at bottom show the timings of saccades (orange) and reward (black) relative to display onset (purple). **B,** Search time and number of saccades (cumulative) for detecting Good object, shown separately for different training amounts (left and right, respectively). Dotted lines: average median across search sessions. Detection of Good objects shows significant increase and median search time and saccade number shows significant decrease by longer reward training. **C,** Distribution of the first saccade directions toward 9 equally spaced objects relative to Good object for different training amounts. Dotted line: chance level. First saccade toward Good object was already higher than chance even after 1-day training and significantly increased by longer reward training. Data in **B-C** are from all four monkeys.

To study the search performance systematically, we used two parameters: 1) reward learning duration (1 day, 5 days, or 30+ days) (Figure 1D) and 2) search display size (DS: 3, 5, 7, or 9 objects) (Figure 1C). For the three learning durations, we used three separate groups of fractals (1-day, 5-day, 30+ day training groups, Figure 1A). None of the fractal groups were used in search task prior to testing. Overall, each monkey learned the values of 212 objects that were later tested in the search task. To test whether the learning effects were retained for a long time, we retested the search performance of the 30+ day group after one month (Figure 1D, memory period). During the memory period, monkeys never viewed any fractals in the 30+ day group, but continued to be engaged in the object-value learning with a new group of fractals (n=72, not used in search).

We found that the object-finding ability improved in two aspects concurrently: accuracy and speed. This learning effect was especially prominent when more objects were present (DS=9), as shown in Figure 2. The average rate of finding the Good object (reflecting ‘accuracy’) grew significantly with more object-reward learning sessions: 71% (1 day), 78% (5 day), and 92% (30+ day) (F_3,44_= 8.4, P=1.5x10^-4^, Figure 2B left). Figure 2B also shows that the average median time to find the Good object (reflecting ‘speed’) across search sessions decreased significantly: 491ms (1 day), 374ms (5 day), and 326ms (30+ day) (F_3,44_=11, P=1.5x10^-5^). Such reduction in search times was accompanied with a reduction in the average median number of saccades to find the Good object across search sessions (F_3,44_=9.6, P=5.4x10^-5^, Figure 2B right). These improvements in the accuracy and speed were maintained after the 1 month memory period with no observable decrement in the 30+ day group (92% detection rate P=0.93, 324ms search time P=0.79).

A typical way to find the Good object was to explore the presented objects by making multiple saccades (Figure 2A). However, after extended reward training, the Good object could be reached directly with a single saccade. This ‘direct choice’ rate significantly improved from 21% in 1-day group to 45% for 30+ day fractal group (F_3,44_=16.7, P=2.1x10^-7^, Figure 2C) and was significantly higher than chance, even for the 1-day group (t_11_=3.6, P= 4x10^-3^). Once more, this direct choice rate was maintained after the 1 month memory period with no observable decrement in the 30+ day group (49% direct choice rate P=0.78). Similar improvements in search speed, accuracy, total saccade number and direct choice rate were observed in other display sizes as reward training duration increased and showed long-term retention for the 30+ day training duration (Figure 3A-C, See **Search Movies**).

**Figure 3.**
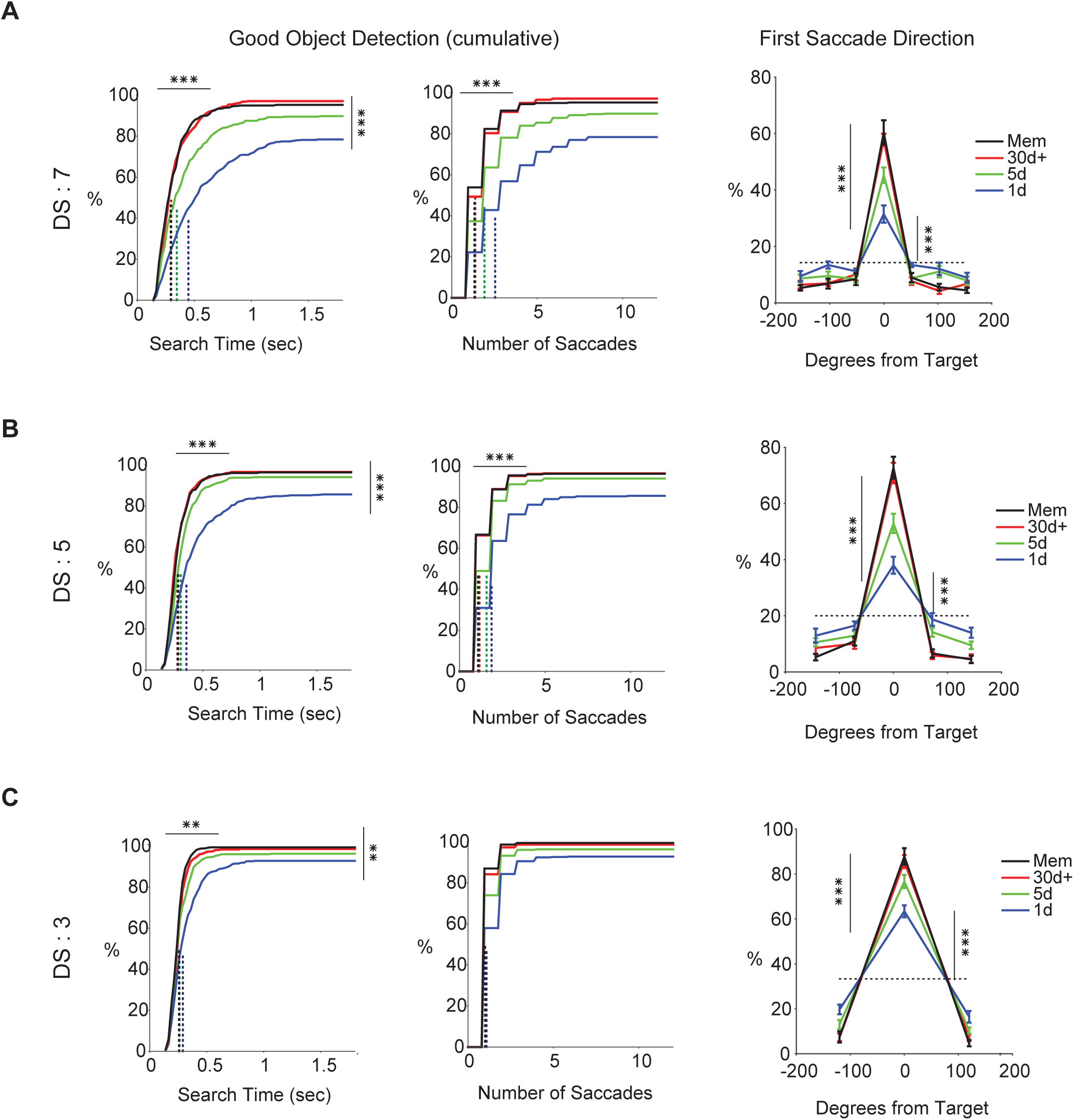
Effects of the repeated object-reward association on the detection of Good objects (DS: 3, 5, 7). **A,** Cumulative distributions of search time (left), the number of saccades (middle) and distribution of the first saccade directions relative to the Good object (right) for DS: 7. Same format as Figure 2B-C. **B-C**, same format as **A** but for DS: 5 in **B** and DS: 3 in **C.** Data are from all four monkeys. Significant increase in detection of Good objects for DS: 3 F_3,44_=4.7 P=5x10^-3^, DS: 5 F_3,44_=6.5 P=9.6x10^-4^, DS: 7 F_3,44_=13.5 P=2.1x10^-6^. Significant decrease in search time for Good objects for DS: 3 F_3,44_=4.3 P=9x10^-3^, DS: 5 F_3,44_=10.8 P=1.8x10^-5^, DS: 7 F_3,44_=10.6 P=2.1x10^-5^. Significant decrease in number of saccades to find Good objects for DS: 5 F_3,44_=11 P=1.5x10^-5^, DS: 7 F _3,44_=10.6 P=2.1x10^-5^ but not in DS: 3 F_3,44_=1 P=0.4. Significant increase in first saccade direction for DS: 3 F_3,44_=12.8 P=3.6x10^-6^, DS: 5 F_3,44_=21.7 P=8.4x10^-9^, DS: 7 F_3,44_=17.5 P=1.3x10^-7^. Significant difference from chance 1-day group for DS: 3 tn=11.5, P= 1.8x10^-7^, DS: 5 t_n_=5.9, P= 9.1x10^-5^, DS: 7 t_n_=5.6 P= 1.5x10^-4^.

Importantly, the direct choice of the Good object started quickly after object presentation. As shown in Figure 4, the first saccade was more likely to be directed to the Good object than a Bad object over all saccade latencies, even when the saccade occurred very quickly (< 150ms). The preferential saccade to the Good rather than a Bad object was enhanced (Figure 4A-B) and appeared to start earlier with longer reward training (significant difference as early as 135ms for 30+ day group tested immediately or after the memory period in search task compared to 175ms for 1-day group, Figure 4C).

**Figure 4.**
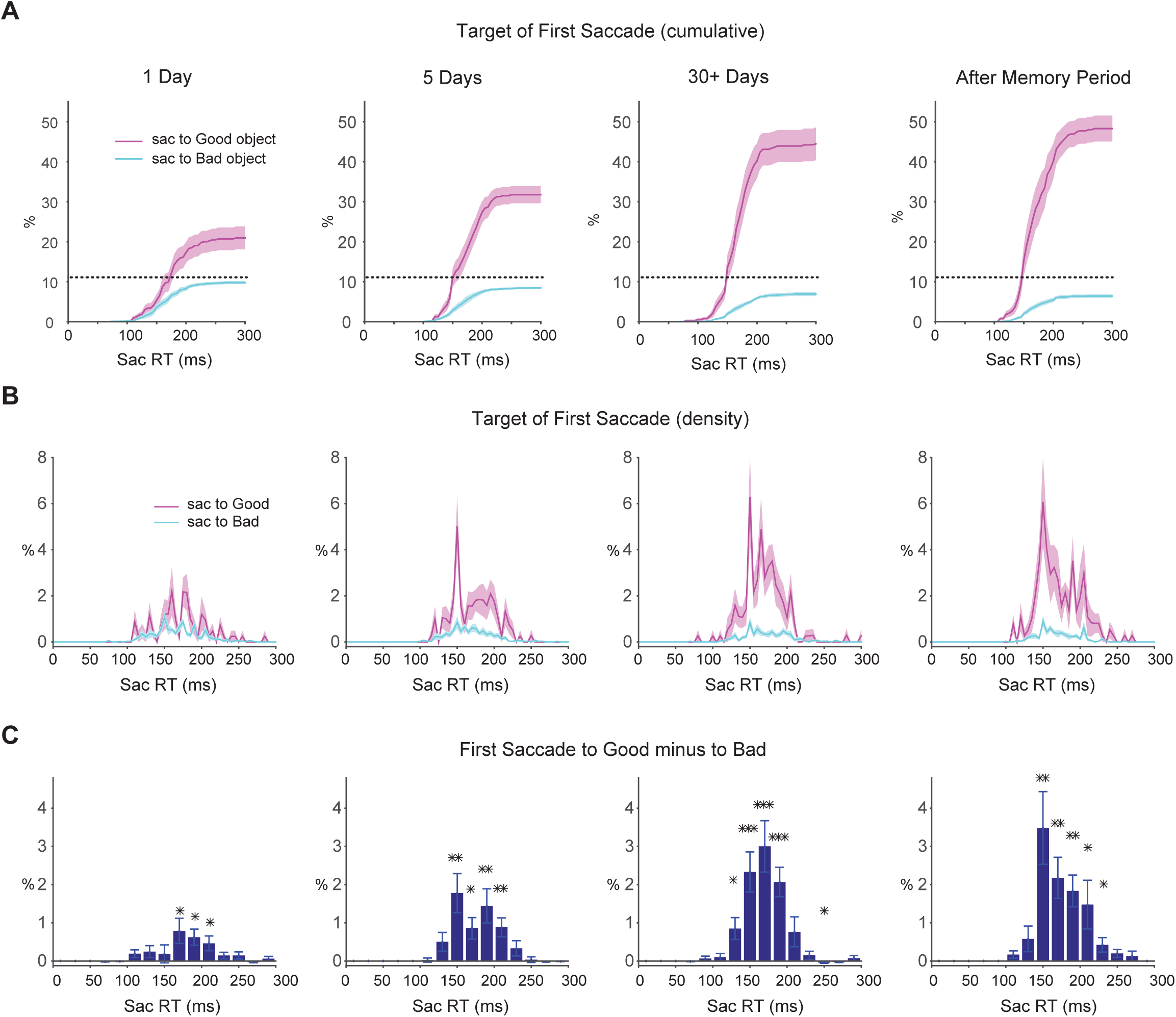
The first saccade bias toward Good objects as a function of saccade reaction time (DS: 9). **A,** Cumulative percentage of the first saccade to Good object (pink) and Bad object (blue) as a function of saccade reaction time. Dotted line: chance level. Preference for the target is evident even at short saccade latencies (<150 ms). Shading shows s.e.m. **B,** Same as **A** but showing differential percentage of the first saccade to Good object (pink) and Bad object (blue). **C,** Saccade percentage to Good minus Bad, binned over saccade reaction time. Paired t-test used for significance in each time bin (two-sided).

Efficient search is characterized by diminishing influence of display size on search time (Wolfe, 1994). Indeed, our results showed a significant interaction between display size and learning duration on search time (i.e., time before reaching Good object) (F_9,173_=3.5, P=5.8x10^-4^, Figure 5A). Analysis of search slopes revealed a significant reduction (~40ms/item to ~16ms/item, F_3,41_=12.8, P=4.78x10^-6^) with longer reward training, consistent with enhanced search efficiency (Figure 5B). Individual subject performance further confirmed consistent and concurrent improvement in search times and search slopes in all participants (Figure 5C).

**Figure 5.**
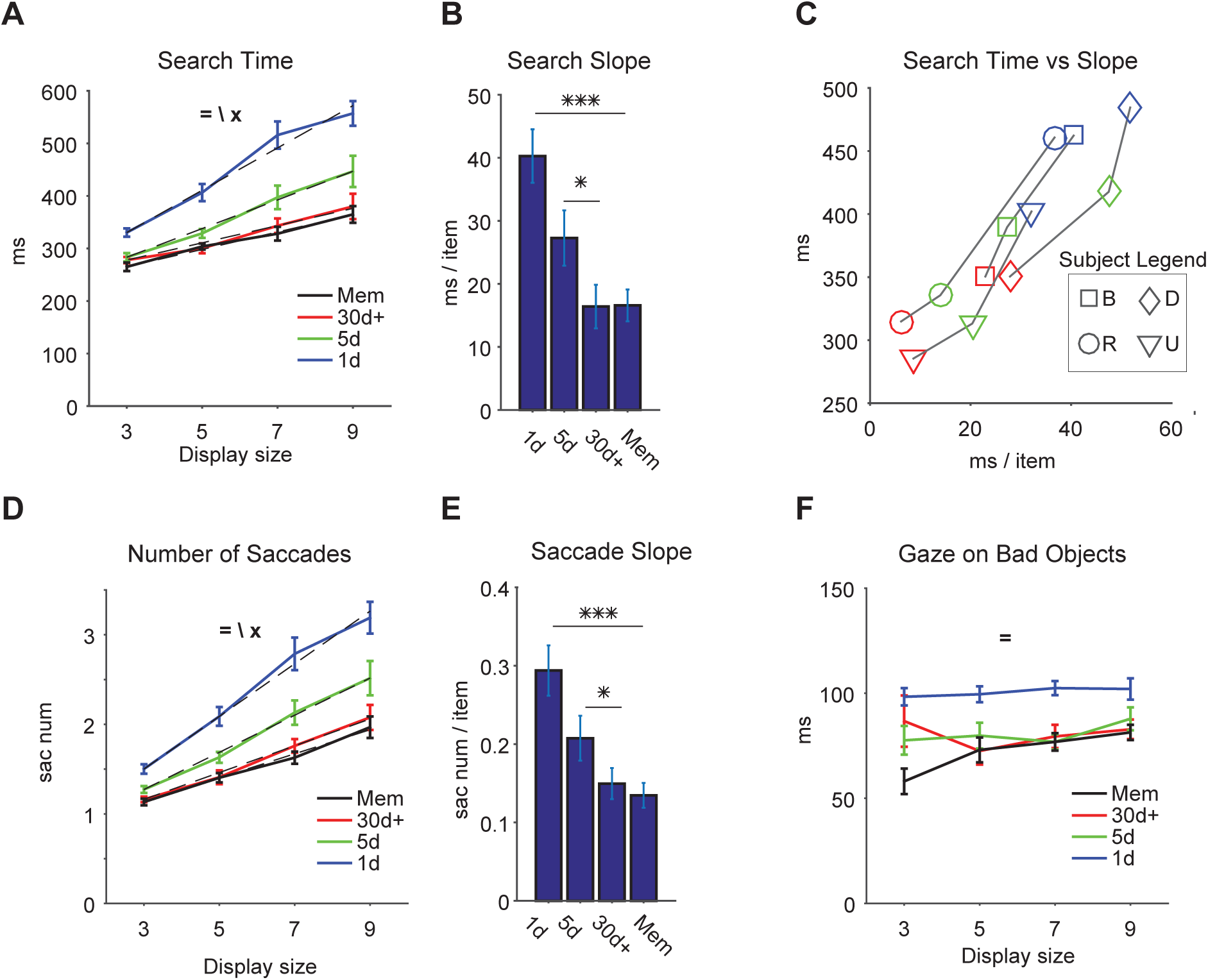
Effects of the display size (DS) on the detection of Good objects. **A,** Time to find the Good object (search time, ordinate) against the number of presented objects (DS, abscissa) for different training amounts and for memory. Dashed lines: linear fits. **B,** Changes in search slope by training amounts and for memory. Post hoc tests show significant change in slope between groups. **C,** Search time (ordinate) and search slope (abscissa) for individual subjects (monkeys B, D, R, U) after different training amounts (blue: 1d, green: 5d, red: 30d+, Mem not shown). **D-E** Number of saccades to find the Good object, shown in the same format as in **A-B. F,** Duration of gaze on Bad objects for different training amounts. (=, \, x: main effect of training amount, display size, and interaction, respectively).

The number of saccades showed a similar trend to search times with saccade slopes reducing as a result of longer reward training (interaction between display size and learning duration F_9,173_=3.5, P=5.5x10^-4^, decrease in slope F_3,41_=11.4, P=1.4x10^-5^, Figure 5D-E). The similarity of trends in search time and number of saccades is consistent with an overt scanning strategy during search when free gaze is allowed as suggested by previous reports (Zelinsky and Sheinberg, 1997; Motter and Belky, 1998). Notably, part of the efficiency in search times came from shorter gaze duration over non-target objects (Bad objects) after longer reward training (Figure 5F). The gaze over Bad objects was ~80ms after extended reward training, which was significantly reduced from ~100ms in 1-day group (F_3,171_=23 P=1.36x10^-12^). Such short inter-saccade intervals is suggested to arise from ‘parallel programing’ of saccades (McPeek et al., 2000).

Importantly, the gained search efficiency turned out to be resistant to the passage of time and interference from new object-reward learning that took place during the 1 month memory period. This can be verified by examining the search time slopes and saccade slopes for the 30+ day group tested immediately or after the 1 month memory period in search task (Figure 5B time slopes: 30d+ 16.4ms/item vs Mem 16.6ms/item and Figure 5E saccade slopes: 30d+ 0.15sacnum/item vs Mem 0.13sacnum/item, P>0.95). These results further confirm the near perfect retention of object finding skill following the 1 month memory period using measures such as search time and direct choice rate described previously (Figure 2-4).

We note that the impaired search performance in 1-day group was not due to incomplete knowledge of object values. This was confirmed by asking the monkeys to make a choice between diametrically presented Good and Bad objects using choice trials (see Methods). Subjects were able to choose the Good objects almost perfectly (>97% in all 4 subjects) with a single session of reward training (n=72 fractals/subject). Thus, search efficiency requires repeated and extended object-reward association beyond what is normally required for making simple value-driven object choice.

## Discussion

Our results demonstrate that finding complex objects in the periphery becomes accurate and quick following long-term reward experience (Figure 2B-C, 3). Search becomes more efficient or less affected by the display size (Figure 5A-E). The Good object is targeted more often by the very first saccade, as if popping out from surrounding objects (Figure 2C).

Studies of visual search and attention have identified a series of visual guiding features (e.g. color) that can aid in the discrimination of a target object from its surrounding and create an easy search (Treisman and Gelade, 1980; Wolfe, 1994; Itti et al., 1998; Itti and Koch, 2000). However, it has been unclear whether non-visual properties of objects can efficiently guide visual search (Wolfe and Horowitz, 2004). Our findings suggest that the long-term object value acts similar to a visual guiding feature with a clear influence on early evoked saccades, consistent with an automatic pre-attentive mechanism (Treisman and Gelade, 1980) (Figure 4).

Our data also suggest important features of the underlying memory mechanism. Monkeys retained the object-finding skill for a long time (long-term memory) for many reward associated objects (high-capacity memory). Importantly, this long-term memory seemed to be unaffected by continual reward learning with novel objects during the memory period. These memory features would be crucial because so many objects are often experienced throughout life and have to be detected in later encounters.

Recent studies suggest that the posterior basal ganglia (Yasuda et al., 2012; Hikosaka et al., 2014), including caudate tail (Kim and Hikosaka, 2013) together with its dopaminergic input from the caudo-lateral substantia nigra (Kim et al., 2014; Kim et al., 2015), contribute to the oculomotor capture to Good objects by mediating long-term object value memories. The output of the posterior basal ganglia, lateral substantia nigra reticulata (SNrl), is shown to exert powerful inhibitory gating on superior colliculus (SC)(Hikosaka and Wurtz, 1983). Importantly SNrl shows rapid and robust inhibition to Good objects. This results in disinhibition of SC and promotion of attention/gaze to the Good objects. On the other hand, when presented with Bad objects, SNrl neurons show strong excitation leading to reduced SC activity and suppression of attention to the Bad object (Yasuda et al., 2012). The differential response of SNrl neurons to Good and Bad objects was found to be virtually unaffected by long memory periods (>100 days), potentially explaining the retention of object finding skill during the 1 month memory period in our task. In addition to SC, the value signals from the posterior basal ganglia circuit may influence cortical areas via thalamo-cortical circuitry (Ilinsky et al., 1985; Deniau and Chevalier, 1992; Middleton and Strick, 2002). Such cortical areas include areas that are involved in visual processing (Middleton and Strick, 1996) and attention (Balan et al., 2008; Cohen et al., 2009), which would contribute to the rapid search for high-valued objects. Future studies will directly examine the neuronal activity in these subcortical and cortical circuits to provide a mechanistic explanation for the emergence of search efficiency observed in the current experiment.

## Acknowledgements

This work was supported by the Intramural Research Program at the National Eye Institute. We thank members of Hikosaka lab for valuable discussions and Simon Hong for providing technical assistance.

## Conflict of interest

Authors do not have any financial or non-financial conflict of interest to declare.

## Materials and Methods

### General procedures

Four adult rhesus monkeys (*Macaca mulatta*) were used for the experiments (monkeys B, R, D male and U female). All animal care and experimental procedures were approved by the National Eye Institute Animal Care and Use Committee and complied with the Public Health Service Policy on the humane care and use of laboratory animals. Monkeys were implanted with a head-post for fixation and scleral search coils to monitor eye movements prior to training in the tasks.

### Stimuli

We created visual stimuli using fractal geometry (Miyashita et al., 1991; Yamamoto et al., 2012). One fractal was composed of four point-symmetrical polygons that were overlaid around a common center such that smaller polygons were positioned more toward the front. The parameters that determined each polygon (size, edges, color, etc.) were chosen randomly. Fractal sizes were on average ~8°x8° but ranged from 5-10 degrees. Each monkey saw three groups of 72 fractals with 1, 5 and 30+ days reward training (1d, 5d and 30d+ training groups, respectively). The order of training groups was randomized between monkeys. Additionally, each monkey saw another group of 72 fractals associated with large or small reward during the 1 month memory period. This group was not used in search task, but was there to ensure that novel reward learning experiences did not interfere with long-term memory results. A separate group of 72 fractals was also used to test monkeys’ value learning after 1-day of training (see **Object-reward training task,** choice trials). Overall, each monkeys saw a total of 360 fractals with biased rewards during this study.

### Behavioral procedures

Behavioral tasks were controlled by custom made C++ based real-time software ‘Blip’ (www.simonhong.org). Data acquisition and output control was performed using National Instruments NI-PCIe 6353. The monkeys sat in a primate chair with their head fixed facing a screen 30cm in front of them. Stimuli generated by an active-matrix liquid crystal display projector (PJ550, ViewSonic) were rear-projected on the screen. Diluted apple juice (33% and 66% for monkeys B, D and monkeys R, U respectively) was used as reward. Rewards amounts could be either small (0.08ml and 0.1ml for monkeys B, D and monkeys R, U respectively) or large (0.21ml and 0.35ml for monkeys B,D and monkey R,U respectively). Eye position was sampled at 1 kHz.

The behavioral procedure consisted of two phases: training (object-reward training task) and testing (search task). Monkeys were trained with separate groups of fractals that differed based on the number of reward training sessions (see **Stimuli**). After the last session of training, performance was tested for 24 fractals per group in each search session. This resulted in three search sessions per group per animal (total of 12 sessions per training group for all four animals).

### Object-reward training task

We used an object-directed saccade task to train object value associations (Figure 1B). Each session of training was performed with a set of eight fractals (4 Good/ 4 Bad fractals). After central fixation on a white dot, one object appeared on the screen at one of the eight peripheral locations (eccentricity 15°). After an overlap period of 400ms, the fixation dot disappeared and the animal was required to make a saccade to the fractal. After 500±100ms of fixating the fractal, a large or small reward was delivered. The displayed fractal was turned off after the time required for large reward delivery for equal perceptual exposure between Good and Bad fractals. This initiated inter-trial intervals (ITI) of 1-1.5s with a blank screen. Each training session consisted of 80 trials with each object pseudo-randomly presented 10 times. Any error resulting from breaking fixation or a premature saccade to fractal resulted in an error tone. Errors were not frequent (<7% of trials). A correct tone was played at conclusion of a correct trial.

The same task structure was used to test choice performance in monkeys with the exception that in choice trials two fractals (1 Good and 1 Bad) were presented diametrically on the screen and monkey was asked to choose one of the fractals after the overlap period by making a saccade to it. Each monkey was trained for 1 day with a new set of 8 fractals (total 9 sessions with 72 fractals per monkey). There were 16 choice trials (randomly selected 1 out of every 5 trials) in an 80 trial training session. The result of choice in the second half of training (last 8 choices) in each session was averaged and used as an estimate of the monkey’s knowledge of object values following 1-day training.

### Search Task procedure

This task started with appearance of a purple fixation dot (Figure 1C). After 400ms of fixation, a display with 3, 5, 7 or 9 fractals were turned on and fixation point was turned off. Fractals were arranged equidistant from each other on an imaginary 15° circle. The location of the first fractal (arbitrary) was uniformly distributed along this circle. Fractals shown in a trial were chosen pseudo-randomly from a set of 24 (12 Good/12 Bad). If gaze left fixation window (5°), the animal had up to 3 seconds to choose an object or reject the trial. Animal could reject a trial either by staying at center (for 600ms) or coming back to center if he did not find a Good object (fixation dot turned back on once gaze left fixation). Next trial would start after 400ms of rejection. Animal was free to make multiple saccades to objects before committing to one by fixating it for at least 400ms (committing time). After committing, animal was required to continue looking for another 100ms (total gaze duration 500ms) after which display was turned off and reward corresponding to the chosen object was delivered. Following reward receipt, an ITI equal to 1-1.5s would ensue. Search task consisted of target present and target absent trials that were intermixed with equal probability. A single Good object was present in target present trials while all objects were Bad in target absent trials (not analyzed in this study). A session of search task consisted of 240 correct trials. Errors included fixation break before display onset (not used in analysis) or fixation breaks after committing to an object (used in analysis). Animals almost never failed to reach a decision (detect or reject) within the 3-second window (%0.08 failed trials).

### Data analysis

Data analysis, plotting and statistical tests were done using Matlab 2014b (MathWorks) using custom written software. Gaze locations were analyzed with an automated script and saccades (displacement>0.5°, peak velocity>50°/s) vs stationary periods were separated in a given trial. Objects were considered to be fixated when gaze was stationary and was within a 5 degree window of their center. Target present trials with rejection, committing to a Bad object or breaking fixation after committing to a Good object were considered misses. Trials with successful gaze to Good objects until reward collection were considered hits. Detection rate (Figure 2B) was the ratio of hits/(hits+misses). For first saccade direction percentage, saccade to each direction (Figure 2C) or object (Figure 4) is divided by the total number of trials with saccades. Slopes of linear fits to search time (Figure 5A-B) and saccade number (Figure 5D-E) were obtained using the Matlab ‘regress’ function.

### Statistical test and significance levels

One-way analysis of variance with training amount was performed for changes in detection rate, median search time and first saccade toward a Good object (Figure 2B-C). Three-way ANOVA with Training x Display size x Subject was performed for search time and saccade number in search task (Figure 5A, D). For effect of training on slopes, two-way ANOVA with Training x Subject was done. Significant ANOVAs were followed by HSD (Honestly Significant Difference) post hoc test when needed (Figure 5B, E). Assumption of normality was confirmed using Lilliefors test in > 94% of cases. Error-bars show standard error of the mean (s.e.m). Significance threshold and marking convention was *p<0.05, **p<0.01, ***p<0.001 (two-sided).

